# Characterization of mycobiota in faba beans infected with *Alternaria* spp

**DOI:** 10.64898/2026.03.19.712847

**Authors:** Biruta Bankina, Ņikita Fomins, Dita Gudrā, Jānis Kaņeps, Gunita Bimšteine, Ance Roga, Frederick Stoddard, Dāvids Fridmanis

## Abstract

Leaf diseases pose a serious threat to faba bean production. Leaf blotch of faba bean, caused by *Alternaria* spp., has become increasingly widespread and destructive in several countries. Leaf diseases pose a serious threat to faba bean production. The infection of plant by pathogens can be influenced by various factors associated with the host plant, environmental conditions and presence of other microorganisms. The phyllosphere and endosphere play a critical role in plant health and disease development. This study aimed to evaluate the factors shaping the structure and diversity of fungal communities associated with faba beans. Plant samples were collected in 2004 from two intensively managed faba bean production fields in the central region of Latvia. Fungal assemblages were characterized using an ITS region metabarcoding approach based on Illumina MiSeq sequencing. Among the assigned amplicon sequence variant (AVS), 65% belonged to the phylum Ascomycota, while approximately 4% were classified as Basidiomycota. *Alternaria* and *Cladosporium* were the dominant genera across samples. The alfa and beta diversities of fungal communities was higher during flowering of faba beans to compare with ripening. The higher abundance of Basidiomycota yeasts were observed during flowering, in contrast, *Cladosporium* genus was significantly more abundant during ripening. *Alternaria* DNA was found on leaves that showed no symptoms of the disease. The diversity and composition of fungal communities were significantly influenced by sampling time and presence of leaf blotch, caused by *Alternaria* spp.

## Introduction

Grain legumes provide protein-rich food and feed, thereby reducing Europe’s dependence on imported proteins. They have considerable potential to enhance cropping systems through biological nitrogen fixation, diversified crop rotations, and other ecosystem services. Faba bean (*Vicia faba* L.) is one of the most important legume crops (Watson *et al*., 2017; Loon *et al*., 2023), and ranks as the fourth most widely cultivated grain legume, with particular significance in the Middle East and Europe (Torabian *et al*., 2024).

Leaf diseases pose a serious threat to faba bean production. In cool-season grain legumes, the principal pathogens include fungi from the genera *Botrytis, Didymella*, and *Uromyces*, as well as genus *Peronospora*, which belongs to the Chromista kingdom. Leaf blotch, caused by *Alternaria* and *Stemphylium*, has become increasingly widespread and devastating in several countries (Bainard *et al*., 2017; Bankina *et al*., 2021; Bruce *et al*., 2025; Ertoy, 2023; Kaur *et al*., 2025; Vaghefi *et al*., 2020). *Alternaria* mycotoxins, restricted under European Union regulations, were isolated from Finnish faba bean seeds in 2024 (Haapalainen, Rämö & Latvala, 2026) further emphasizing the importance of understanding this disease. The symptoms induced by *Alternaria* spp. and *Stemphylium* spp. are highly similar and, in many cases, indistinguishable; therefore, reliable identification under field conditions is not possible. In addition, there is no confirmed information regarding the specific species within these genera that are involved in the disease. For these reasons, the designation *Alternaria/Stemphylium* is more appropriate for referring to the causal agents of this disease. Overall, studies examining this disease and the factors influencing its development and severity remain limited.

The infection of plant by pathogens can be influenced by various factors associated with the host plant, environmental conditions and presence of other microorganisms. Numerous studies have focused on fungal communities in soil and the rhizosphere (Gao *et al*., 2021; Granzow *et al*., 2017), whereas considerably less attention has been given to the endosphere of aerial plant tissues. To date few studies have addressed the fungal communities associated with faba bean phyllosphere and endosphere (Li *et al*., 2024; Li, Hou & Liu, 2025).

Significant scholarly inquiry has been devoted to characterizing shifts in mycobiota composition and structural organization of microbial communities in response to the presence of plant pathogens (Dai *et al*., 2022; Diáz-Cruz & Cassone, 2022; Tao *et al*., 2021). However, microbial communities are highly diverse and interact differently with various hosts, leading to inconsistent results. Existing findings remain inconsistent and definitive patterns have yet to be identified.

Recent studies have shown that infection by pathogens, including *Alternaria*, significantly changes microbial communities (Tao *et al*., 2021), but there is little information on the what happens to those communities during disease development. Despite growing interest in plant-associated microbiomes, knowledge about the potential role of community-level foliar mycobiota in promoting plant health remains limited. Therefore, the analysis of fungal communities can provide insights into the factors that affect the likelihood of pathogenic infection in plants (Li *et al*., 2024).

The aims of this research were (1) to evaluate the factors shaping the structure and diversity of fungal communities associated with faba bean plants, and (2) determine how fungal community composition differs between plants exhibiting visible symptoms putatively associated with species of the genera *Alternaria*/*Stemphylium* and asymptomatic plants’ parts. Hence, our research question was to investigate the composition of the mycoflora in field conditions. By comparing the composition between leaves with and without leaves, we aimed to test for underlying differences associated with the development of diseases, caused by *Alternaria* spp.

## Materials and Methods

### Sample collection and preparation for subsequent analyses

Samples were collected in 2024 from two intensively managed faba bean production fields in the central region of Latvia (56.57°N, 23.38°E and 56.33°N, 24.20°E), where the cultivar ‘Fuego’ was grown. Plants were sampled three times: during the first decades of June, July, and August, corresponding to the flowering, pod development, and ripening stages (BBCH growth stages 65, 73, and 80, respectively). At each sampling date and location, ten plants were randomly selected and cut off at the soil surface.

Immediately after collection, plants were separated into leaves, stems, flowers, and pods with seeds. Plant parts were categorized as either asymptomatic or showing symptoms characteristic of *Alternaria*/*Stemphylium* infection.

Samples were rinsed three times with autoclaved distilled water, blotted dry between sterile sheets of filter paper, and placed in plastic bags. All samples were stored at −80°C until molecular analyses were performed. A total of 186 plant samples were collected across all sampling dates and locations: 90 in June, 72 in July, and 24 in August, with the lower number in August attributable to plant senescence. Among the samples, 81 consisted of leaves, 40 of stems, 40 of pods with seeds, and 25 of flowers; 85 exhibited visible symptoms putatively caused by *Alternaria* and/or *Stemphylium*, whereas 101 showed no visible symptoms.

### Identification of causal agents of diseases

Each symptomatic sample was sectioned into two parts: one part was preserved by freezing as previously described, and the other part was processed for pathogen isolation into pure culture.

Small tissue segments from infected tissues were surface-sterilized in 1% sodium hypochlorite for 1 min, rinsed three times in sterile distilled water, and plated onto potato dextrose agar (PDA) supplemented with streptomycin (100 mg L^−1^) and penicillin (100 mg L^−1^). Single-hyphal-tip isolates were recovered from each emerging colony and sub-cultured on PDA to obtain pure cultures. The isolates were first characterized based on morphological features and subsequently subjected to molecular analysis of the ITS region using the ITS1-F and ITS4 (reverse) primers, as previously described (Brauna-Morževska *et al*., 2023). Morphological characteristics and molecular analyses confirmed the preliminary visual identification – the visible symptoms were caused by *Alternaria* and/or *Stemphylium* spp.

### Sample preparation for high throughput ITS sequencing

For DNA extraction, 150 mg of plant tissue was transferred to a Lysing Matrix A tube (MP Biomedicals, USA), which was prefilled with lysing buffer that was supplemented with 1.5% polyvinylpyrrolidone (PVP) (Sigma-Aldrich, USA) and processed using the FastDNA Spin Kit for Soil (MP Biomedicals, USA) according to the manufacturer’s instructions. The concentration of the extracted DNA was determined using a Qubit dsDNA HS Assay Kit and a Qubit Flex Fluorometer (Thermo Fisher Scientific, USA). The choice of extraction procedure was based on our previous experience with fungal DNA extraction from weed seeds (Nečajeva *et al*., 2024).

A two-stage PCR protocol was used for MiSeq library preparation. Primers containing Illumina overhang adapters were designed for PCR amplification of the fungal ITS gene (Table 1). Fifty nanograms of DNA were amplified in triplicate using fungal primers targeting the fungal ITS gene. PCR reactions were performed in a total volume of 20 µL using the Phire Plant Direct PCR Kit (Thermo Fisher Scientific, USA), which tolerates higher levels of PCR inhibiting adducts, under the following conditions: initial denaturation at 98°C for 5 min; 40 cycles of denaturation at 98°C for 5 s, annealing at 65°C for 5 s, (initially calculated employing Thermo Fisher Scientific “Tm Calculator for Primers” and the optimized to identify conditions that generate PCR product at highest yield and specificity, and extension at 72°C for 20 s; followed by a final extension step at 72°C for 1 min. To ensure the purity and reliability of the PCR products, negative controls were included for each prepared master mix. The yield of PCR products was assessed by 1.2% agarose gel electrophoresis, and the products were purified using the NucleoMag NGS Clean-Up and Size Select kit (Macherey-Nagel, Germany). PCR product concentrations were measured using a Qubit dsDNA HS Assay Kit and a Qubit Flex Fluorometer (Thermo Fisher Scientific, USA). Amplicons from each replicate were pooled in equal quantities, and samples were normalised to 5 ng/µL. During the second PCR stage, Illumina MiSeq i7 and i5 indices were added to 5 ng of ITS PCR product using custom-ordered Nextera XT Index Kit primers (Illumina Inc., USA; Metabion International AG, Germany). Before sequencing, all samples, including negative controls, were pooled at equimolar concentrations and diluted to 6 pM. Sequencing was performed using paired-end chemistry with a 500-cycle MiSeq Reagent Kit v2 on an Illumina MiSeq platform (Illumina Inc., USA), targeting a minimum sequencing depth of 20,000 reads per sample.

**Table 1.**
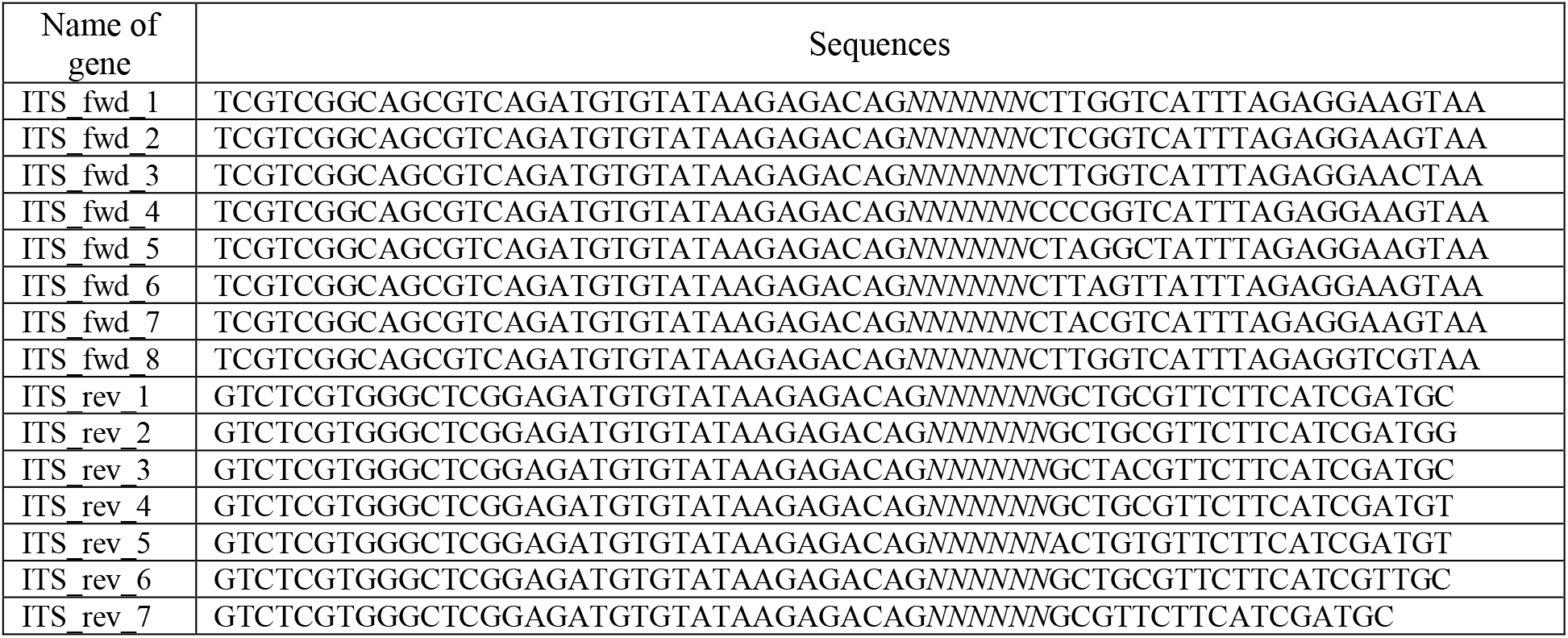
The primers used in this study with Illumina overhang adapters to amplify the fungal ITS gene.

### ITS sequence analysis

Sequence reads were quality filtered and trimmed using fastp with the following parameters: -f 30 -F 30 - 5 20 -3 20 -q 20 -l 40 --correction. All quality-approved reads were imported into QIIME2 v.2024.2 (Bolyen *et al*., 2019) for downstream analysis. The DADA2 (Callahan *et al*., 2016) plugin was used for paired-end read merging, quality control, and denoising; phiX reads and chimeric sequences were removed using the pooled consensus method. Multiple sequence alignment was performed *de novo* using the MAFFT (Katoh & Standley, 2013), and a phylogenetic tree was constructed with FastTree2 (Price, Dehal & Arkin, 2010). Taxonomic assignment was carried out using the UNITE fungal ITS database v.10 (dynamic release; downloaded September 2025) (Nilsson *et al*., 2019) trained with a naïve Bayes classifier (Pedregosa *et al*., 2011).

Statistical analyses and figure generation were conducted using R version 4.3.2. Raw reads were initially filtered to remove low-abundance and infrequently detected taxa using the subset by prevalent function from the mia package (detection = 10/1000, prevalence = 5/100) (Borman *et al*., 2025). The resulting dataset was rarefied to a sequencing depth of 7,000 reads.

Alpha diversity (Shannon and inverse Simpson indices) was calculated at the genus level using mia v1.17.9. Group differences were evaluated using the Mann–Whitney U test and the Kruskal–Wallis test, with statistical significance defined as p ≤ 0.05.

Beta diversity was assessed using Bray–Curtis dissimilarities computed with phyloseq v1.53.0 (McMurdien & Holmes, 2013) and visualized via principal component analysis (PCA). Community-level differences were tested using permutational multivariate analysis of variance (PERMANOVA), and homogeneity of group dispersions (PERMDISP) was evaluated using vegan v2.6-4. Multilevel pairwise comparisons were performed with pairwise A-donis.

Differential abundance of fungal genera between groups was analyzed using ANCOMBC2 (Lin & Peddada, 2024), with significance defined as padj.FDR < 0.05.

Differential network analysis was conducted using NetCoMi v1.1.0 (Peschel *et al*., 2021). Correlations were computed using Spearman’s method. Associations between taxa were evaluated using t-tests, with a significance threshold of 0.05.

Data visualization was performed using ggplot2 v4.0.0 (Wickham, 2016).

## Results

### Overview of fungal taxa detected in faba bean samples

A total of 6,129,116 high-quality ITS reads were obtained from 186 faba bean samples. The raw dataset comprised 915 amplicon sequence variants (ASVs) representing 194 genera. Communities exhibited extreme taxonomic unevenness, with pronounced dominance by a limited number of taxa.

Of the assigned ASVs, 65% belonged to the phylum Ascomycota, and about 4% were classified as Basidiomycota. The remaining ASVs were affiliated with Chytridiomycota, Mucoromycota, *Incertae sedis* taxa, or could not be assigned due to the absence of significant sequence similarity in reference databases.

At the order level, fungal communities were strongly dominated by *Cladosporiales* (45%), followed by *Pleosporales* (19%) within the phylum Ascomycota, and *Tremellales* (3%) within Basidiomycota.

*Cladosporium* was the most abundant genus (44.9% of all reads), followed by unclassified fungi (30.8%) and *Alternaria* (18.5%), collectively accounting for approximately 95% of the total fungal community (Figure 1). The remaining genera constituted a substantial rare biosphere, each typically representing <0.5% of total reads and often detected sporadically in only one or two samples, suggesting transient presence rather than stable community membership.

**Figure 1.**
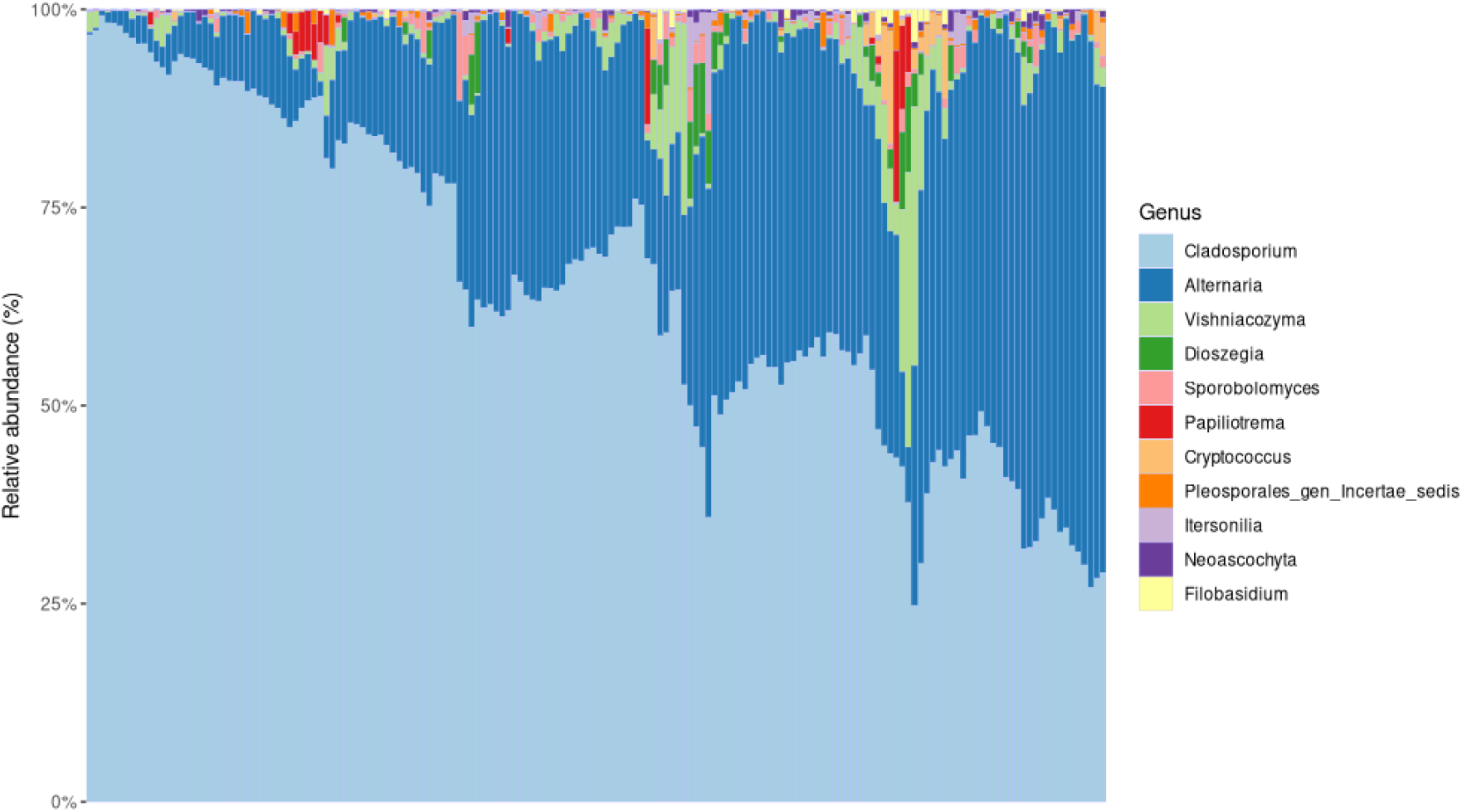
Relative abundance of fungal genera depending on samples.

Rarefaction, and removal of low-abundance and unclassified taxa yielded a total of 11 fungal genera across all samples (Table 2.) Per-sample fungal ASV richness ranged from 3 to 11 (median = 7). The refined fungal community was dominated by *Cladosporium* (mean relative abundance = 65.6%) and *Alternaria* (28.1%), which were the most abundant classified genera. However, both taxa exhibited substantial inter-sample variability, as reflected by high standard deviations (20.3% and 18.3%, respectively). Given that the relative abundance of *Stemphylium* was < 0.001, *Alternaria* was considered the causal agent of leaf blotch in 2024.

**Table 2.** Mean relative abundance and variability of fungal genera in faba bean samples.

| No | Genus | Phylum | Relative abundance (proportion) |  |
| --- | --- | --- | --- | --- |
|  |  |  | Mean | ± SD |
| 1 | *Cladosporium* | Ascomycota | 0.656 | 0.203 |
| 2 | *Alternaria* | Ascomycota | 0.281 | 0.183 |
| 3 | *Vishniacozyma* | Basidiomycota | 0.024 | 0.049 |
| 4 | *Dioszegia* | Basidiomycota | 0.008 | 0.022 |
| 5 | *Sporobolomyces* | Basidiomycota | 0.007 | 0.013 |
| 6 | *Papiliotrema* | Basidiomycota | 0.005 | 0.021 |
| 7 | *Cryptococcus* | Basidiomycota | 0.004 | 0.017 |
| 8 | *Genus Incertae sedis, Pleosporales* | Ascomycota | 0.004 | 0.008 |
| 9 | *Itersonilia* | Basidiomycota | 0.004 | 0.013 |
| 10 | *Neoascochyta* | Ascomycota | 0.003 | 0.005 |
| 11 | *Filobasidium* | Basidiomycota | 0.002 | 0.006 |

### Differences in fungal communities depending on sampling site, plant organs and sampling time

Fungal communities across different plant organs (leaves, stems, flowers, and pods with seeds) exhibited no significant differences in α-diversity or β-diversity. Likewise, sampling site did not significantly influence fungal community diversity or composition.

In contrast, the composition of phyllosphere and endosphere fungal communities was significantly shaped by sampling time, corresponding to plant developmental stage. The ɑ-diversity varied significantly across months (Figure 2). Both Shannon and inverse Simpson indices showed strong overall differences among flowering (June), development of fruit (July), and ripening (August) (Shannon: Kruskal–Wallis H(2) = 21.733, P < 0.001; inverse Simpson: H(2) = 14.78, P < 0.001). Pairwise tests revealed that diversity was significantly lower in June than in July (Shannon: P = 4.16 × 10^−6^; inverse Simpson: P = 0.0001). Differences between June and August, and between July and August were not statistically significant. These results indicate a pronounced increase in fungal diversity from early to mid-season, followed by stabilization between July and August.

**Figure 2.**
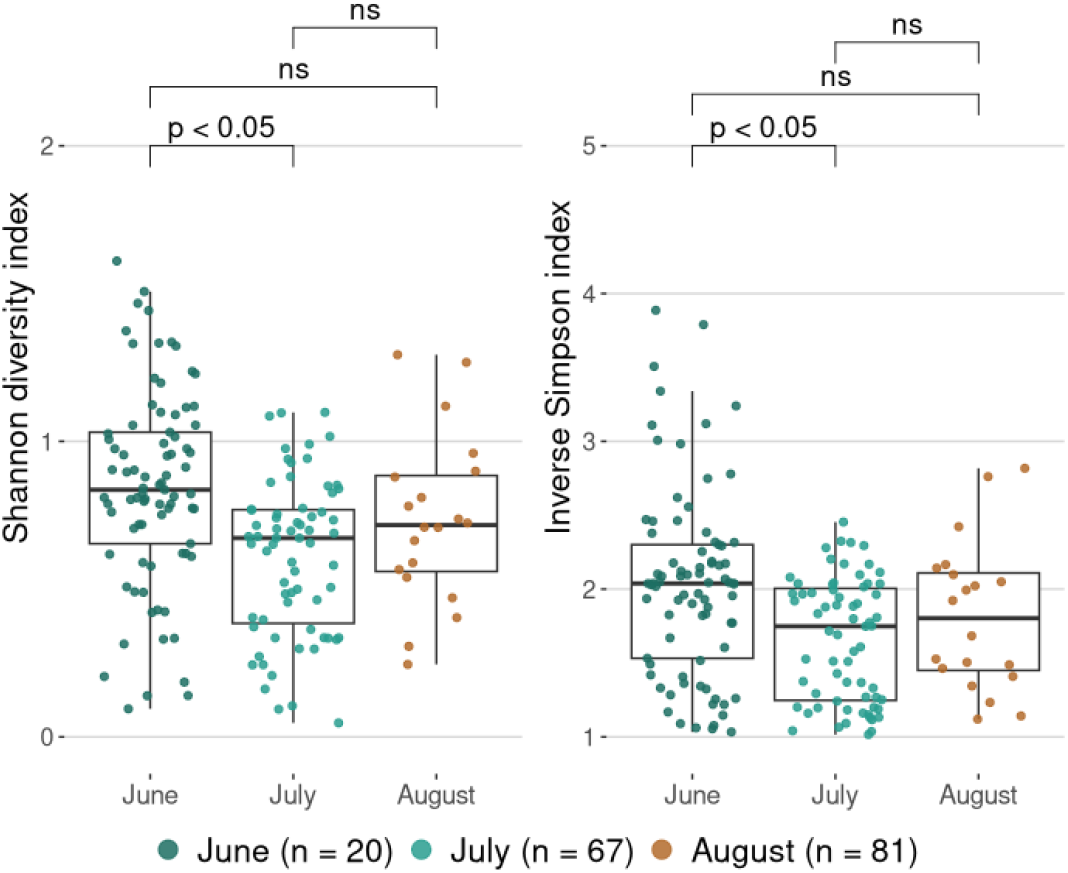
The ɑ-diversity of fungal communities depending on sampling time.

The β-diversity analyses supported this seasonal shift (Figure 3). PERMANOVA revealed month-to-month shifts in community composition (R^2^ = 0.04997, F = 4.339, P = 0.0061), although the effect size was modest. However, a test for homogeneity of multivariate dispersions showed significant differences in within-group variability among months (F = 4.193, P = 0.0154), indicating that part of the PERMANOVA signal may stem from dispersion rather than true compositional differences.

**Figure 3.**
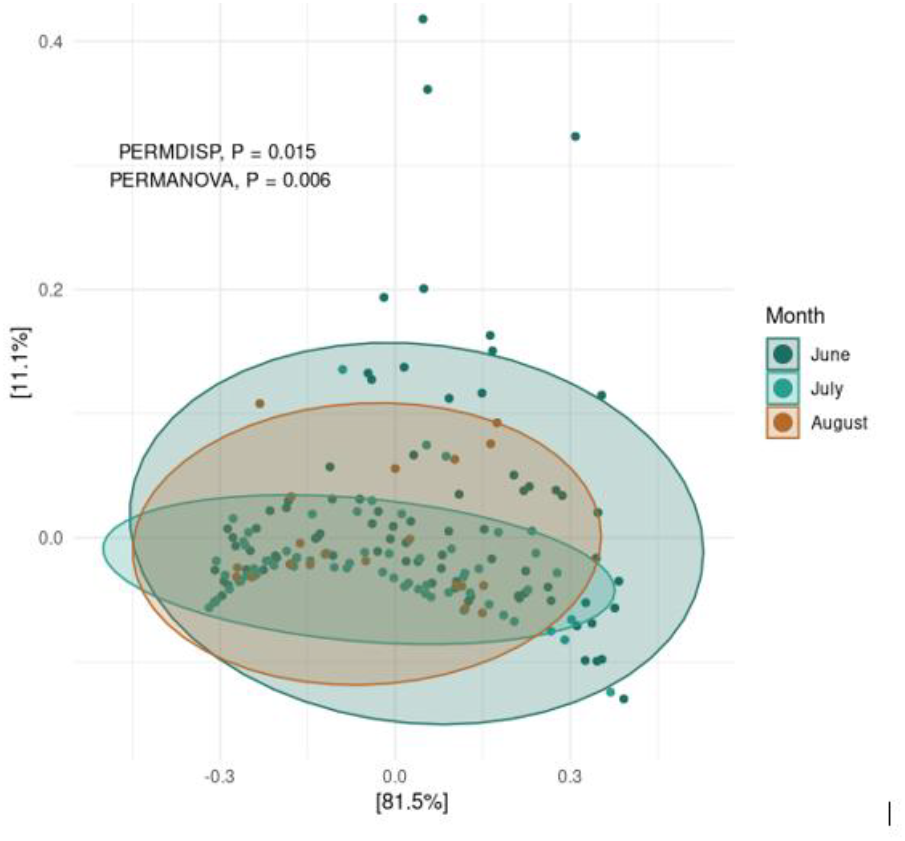
Variation in fungal community β-diversity across sampling times as revealed by principal coordinates analysis (PCoA) using Bray–Curtis dissimilarity.

Differential abundance analysis using ANCOM-BC2 identified several fungal genera that differed significantly between flowering and development of fruit (Figure 4). No significant differences in fungal genus composition were observed between August and the other sampling months. The yeast genus *Sporobolomyces* exhibited the strongest temporal shift, with a substantially higher abundance during flowering than time of fruit development (q = 7.25 × 10^−6^). Other Basidiomycota yeasts, including *Itersonilia, Dioszegia*, and *Vishniacozyma*, were significantly enriched in June (q < 0.05). In contrast, *Cladosporium* was the only genus that was significantly more abundant in July than in June (q = 0.0039). No other genera exhibited statistically significant differences between the two months.

**Figure 4.**
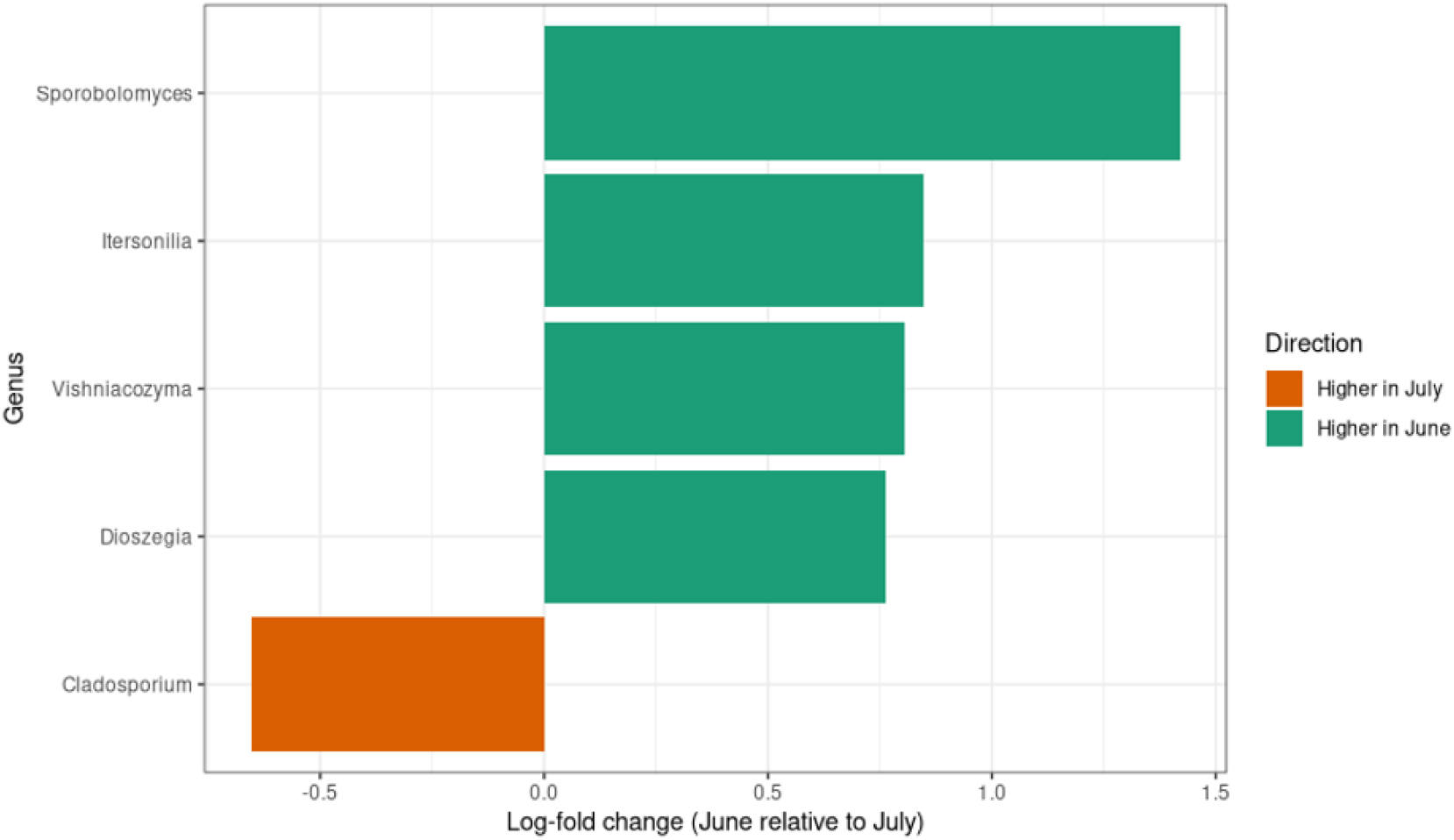
Differentially abundant fungal genera between flowering and fruit development.

Differential network analysis revealed significant differences in fungal co-occurrence only between June and July (Figure 5). June was characterized by strong positive associations among the Basidiomycota yeasts *Vishniacozyma, Filobasidium*, and *Cryptococcus*, as well as a distinct linkage between *Itersonilia* and *Sporobolomyces*, indicating early-season clustering of yeast taxa. In July, these yeast–yeast associations persisted and were further expanded by the integration of *Dioszegia*, reflecting increased connectivity within the yeast community. In contrast, the July network exhibited a structural shift toward filamentous fungi, with *Alternaria* showing stronger associations with other *Pleosporales* taxa, suggesting a mid-season intensification of interactions within *Dothideomycetes*. Several genera remained central across both months, but the overall topology and dominant associations differed, reflecting seasonal restructuring of fungal community interactions.

**Figure 5.**
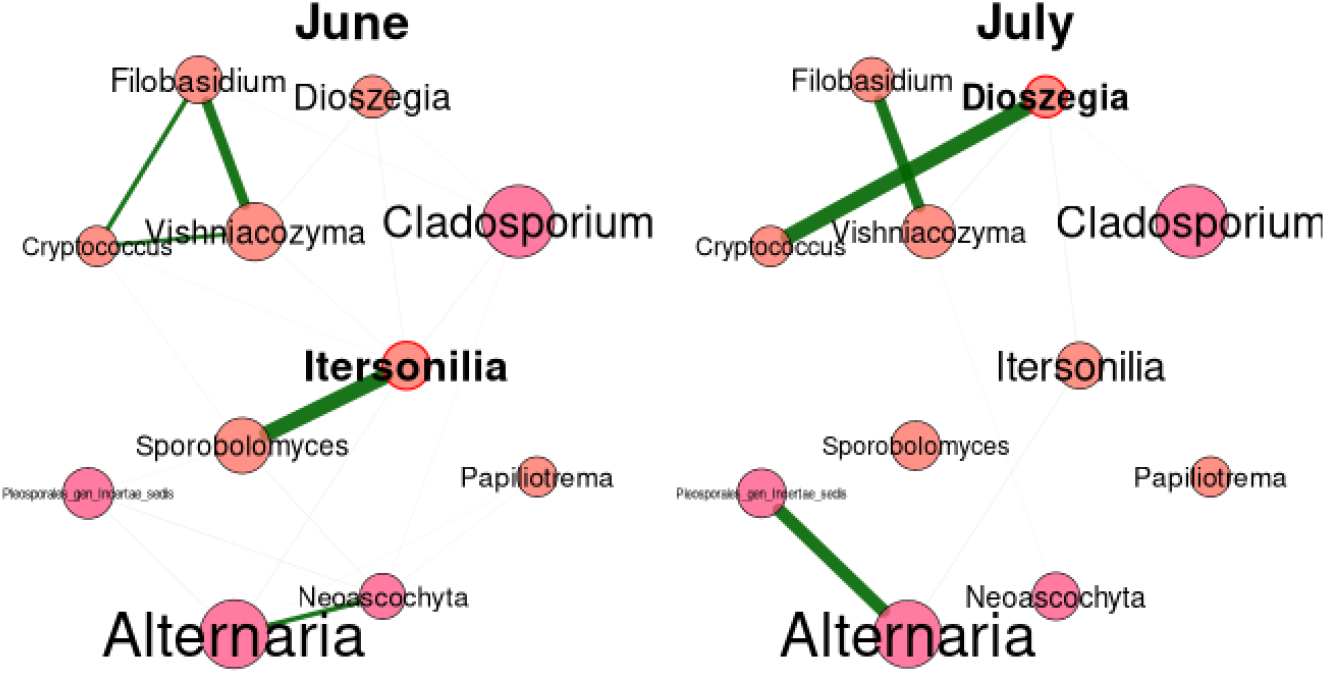
Seasonal variation in fungal co-occurrence networks between flowering and fruit development

### Composition of fungal communities as influenced by presence of visible symptoms of leaf blotch, caused by Alternaria

Fungal community composition was significantly influenced by the presence of visible leaf blotch symptoms caused by *Alternaria* spp. The ɑ-diversity analysis revealed a significant difference between groups with and without *Alternaria* symptoms (Figure 6). Samples exhibiting symptoms had lower diversity, as indicated by the Shannon index (Mann–Whitney U test, W = 4212, P = 0.024) and inverse Simpson’s index (W = 4203, P = 0.026).

**Figure 6.**
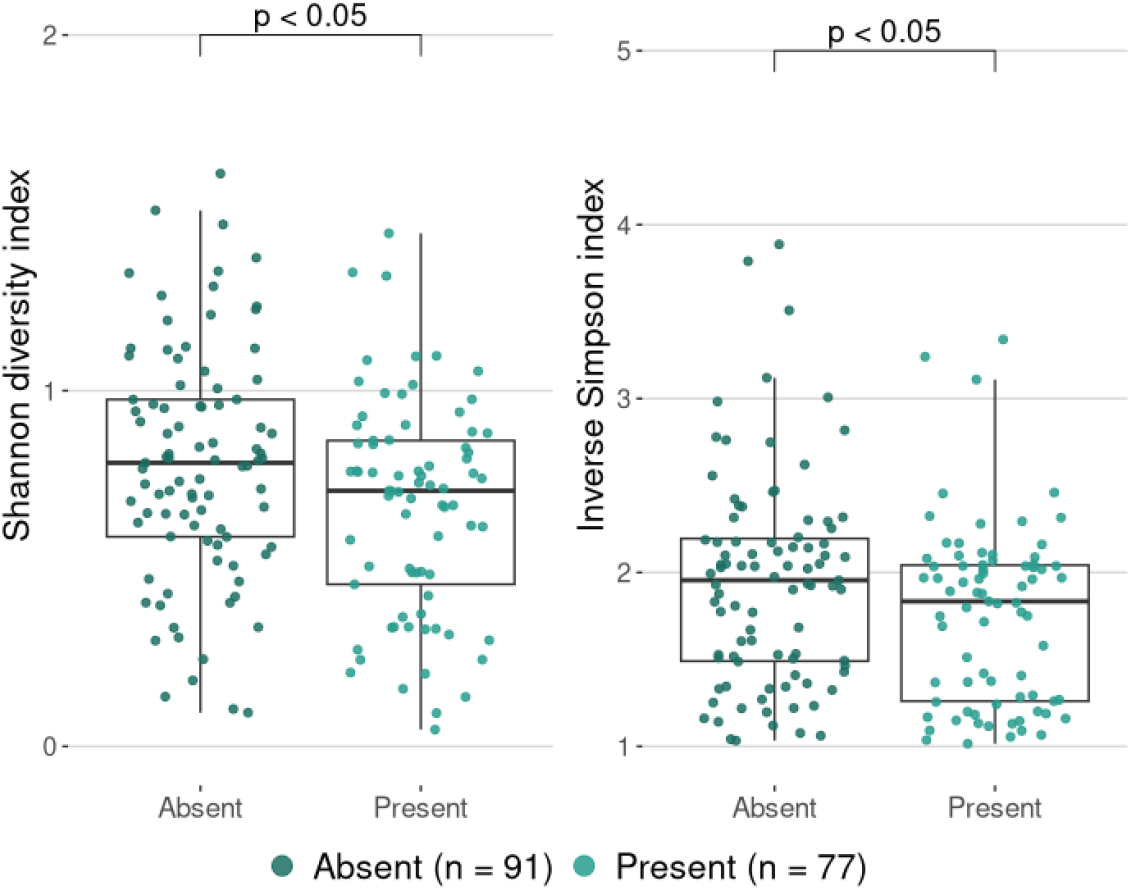
Shannon and inverse Simpson diversity indices in relation to the presence of visible leaf blotch symptoms caused by *Alternaria* spp.

The β-diversity analysis showed no significant differences in overall fungal community composition between samples with and without *Alternaria* symptoms (Figure 7).

**Figure 7.**
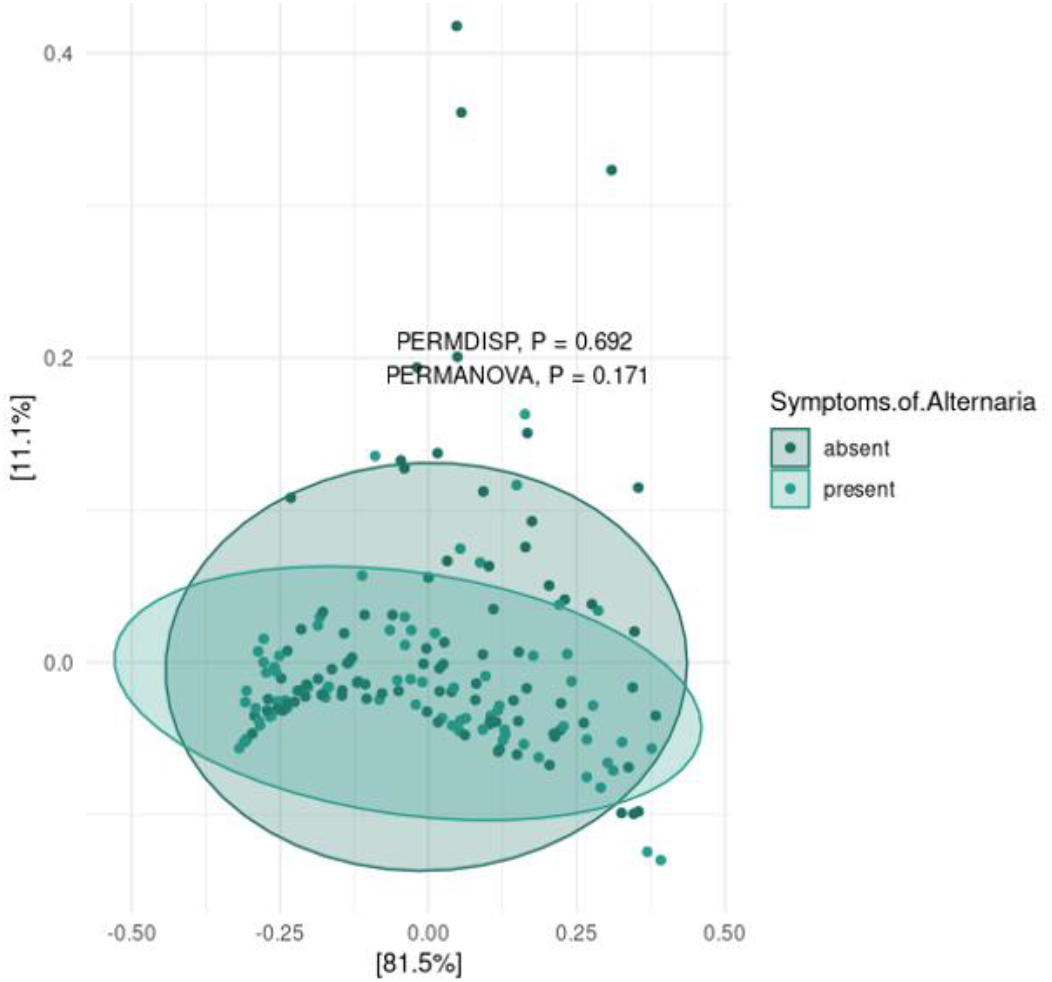
Variation in fungal community β-diversity between symptomatic and asymptomatic samples revealed by Bray–Curtis-based PCoA.

Together, these results suggest that while ɑ-diversity was reduced in symptomatic samples, the presence of *Alternaria* symptoms was not associated with distinct shifts in overall fungal community structure.

## Discussion

We found that the fungal community changed through the growing season in two field of faba bean. *Alternaria* was found on almost all samples, whether or not there were disease symptoms. Fungi residing on and within plant tissues, together with the host, form an interconnected system that is strongly shaped by the environment. These intricate interactions play a critical role in regulating plant physiological processes and overall plant health.

Fungi belonging to the phylum Ascomycota dominated all sample categories examined in this study, with members of Basidiomycota representing the second most abundant group of identified fungi. This community structure is consistent with numerous previous studies across a wide range of plant hosts and countries (Tao *et al*., 2021; Kaniyassery *et al*., 2024;). Notably, similar patterns have been reported in faba bean leaves in Germany, where *Ascomycota* constituted more than 70% of the fungal community (Granzow et al., 2017). Likewise, Li et al. identified Ascomycota and Basidiomycota as the principal components of fungal communities associated with faba beans leaves in China (Li *et al*., 2024; Li, Hou & Liu, 2025).

The predominance of *Ascomycota* is expected, as the phylum includes many saprotrophs inhabiting plants, as well as major faba bean pathogens belonging to the orders *Cladosporiales* and *Pleosporales*, both of which were dominant in the presented study.

*Cladosporiales* is a diverse order, containing species with different life styles – saprotrophic, endophytic, fungicolous, lichenicolous, and pathogenic on humans and plants (Abdollahzadeh *et al*., 2020). The genus *Cladosporium* is often a significant member of fungal communities on different plant leaves (Huang *et al*., 2021; Kaniyassery *et al*., 2024). As found in an investigation of fungal assemblages of faba bean leaves in China (Li *et al*., 2025), *Cladosporium* dominated in the present studies. However, assessing its impact on plant health is difficult, because *Cladosporium* spp. can occupy many ecological niches. They may function as pathogens, saprotrophs, and potentially also as endophytes or epiphytes.

*Alternaria* has been reported as the dominant genus on a variety of plant species, including maize leaves (Luo *et al*., 2023), broccoli (Kim & Park, 2023), and in the endosphere of faba bean in China (Li, Hou, & Liu, 2025). Consistent with these findings, the present study showed that *Alternaria* was the second most widespread fungal genus detected.

The genus *Alternaria* comprises more than 368 species in 29 sections with different trophic modes, including plant pathogens. *Alternaria* spp. have been recognized as important pathogens of faba bean and other legumes, but there are few studies related to the identification of species that are pathogenic to legumes (Baida, Abass, & Salih, 2025; Nichea et al., 2022). Only the ITS region was targeted using a high-throughput sequencing approach; consequently, taxonomic resolution was limited and species-level identification was not achieved. The trophic modes of *Alternaria* within the fungal communities were not analysed, as species within this genus encompass diverse ecological guilds.

*Vishniacozyma, Dioszegia, Sporobolomyces* and other yeasts from Basidiomycota were detected at relatively high abundances, indicating a notable contribution of yeast-forming *Basidiomycota* to the overall fungal community. The genus *Vishniacozyma* has been reported as a dominant taxon of yeasts on faba bean leaves in a study conducted in China (Li, Hou, & Liu, 2025). *Filobasidium, Papiliotrema* and *Sporobolomyces* are typical yeast genera frequently identified on a wide range of plant species (Kim & Park, 2023). Despite their consistent presence and, in some cases, high relative abundance in plant-associated microbiomes, the roles of Basidiomycetous yeasts in plant development, plant health, and their interactions with other microorganisms remain largely unclear.

The structure and diversity of fungal communities were not influenced by either sampling site or plant organ. This pattern may be explained by the strong dominance of the genera *Cladosporium* and *Alternaria*, which likely masked minor differences in the relative abundance of less prevalent taxa. Additionally, the same cultivar ‘Fuego’ was produced in both sites, which could have contributed to the similarity of fungal assemblages.

Only aerial plant parts were analysed in this study, and the stems, leaves, pods, and flowers harboured broadly similar fungal assemblages. These above-ground organs are exposed to comparable environmental conditions, which may promote homogenization of fungal communities across plant parts. Similar results have been reported in relation to aerial organs of cashew, where the lack of differentiation of fungal communities among different organs was noted (Mukhebi *et al*., 2024).

The phyllosphere and endosphere of plants change in line with the age of plants (Shi *et al*., 2024), but results are not consistent. In the present work, the ɑ-diversity of fungal communities was essentially higher in June (GS 65) than in July (GS 73). This could be associated with differences in plant age and development stage. Similar seasonal changes in diversity of fungal communities have also been revealed in other situations (Shi *et al*., 2024).

In general, a higher abundance of yeasts was detected in June, coinciding with the flowering stage of faba bean. Consistent results were obtained in China, with a decline in the relative abundance of *Filobasidium* during tobacco leaf maturation. (Shi *et al*., 2024). The increased prevalence of yeasts during this period may be attributed to the higher availability of readily utilizable nutrients in young aboveground plant tissues. In contrast, the relative abundance of *Cladosporium* increased in July, and many species within this genus exhibit a necrotrophic lifestyle and preferentially colonize senescing or damaged plant tissues.

The severity of leaf blotch, caused by *Alternaria* and *Stemphylium* increases rapidly during the stage of faba bean flowering and early pod development in July under Latvian (Bankina *et al*., 2021) and Canadian conditions (Bruce *et al*., 2025; Kaur *et al*., 2025). Network analyses in the present study clearly show connectivity between *Alternaria* and other *Pleosporales* in July. This allows us to make a hypothesis regarding the complexity of the causal agents of leaf blotches, as observed under field conditions.

In our study, α-diversity (Shannon and Simpson inversion indexes) was higher in plant tissues without symptoms. Similar results were obtained in other research – Shannon indexes clearly demonstrated higher diversity in the phyllosphere and endosphere of healthy faba bean leaves compared with diseased plants in China (Li, Hou & Liu, 2025). Similar tendencies were observed, where diseased leaves showed lower values of these indexes than healthy leaves of both tobacco and tea (Huang *et al*., 2021; Guo *et al*., 2025). However, results are not consistent; diversity of fungal communities changed in correlation with severity of angular spot of cucumbers, but clear tendencies were not determined (Luo *et al*., 2025). Díaz-Cruz and Cassome (2022) observed no significant differences in fungal α-diversity in the phyllosphere of healthy soybean leaves and those infected by *Phytophthora sojae* and *Septoria glycines* leaves. Although notable distinctions in the fungal communities in samples from healthy and diseased faba beans were observed in China (Li, Hou & Liu, 2025), our study did not support these findings. Differences in β-diversity was not observed to depend on presence of visible symptoms in our study.

A clear distinction in mycobiota composition has been detected between healthy and diseased plants elsewhere (Li *et al*., 2024). However, this observation was not confirmed in presented study. It is possible to conclude, that the presence of *Alternaria* was high in all parts of plants, independently of the presence of visible symptoms. The results of this study highlight the need for further investigations, particularly regarding the identification of *Alternaria* species and their pathogenicity to faba beans.

## Conclusion

The genera *Cladosporium* and *Alternaria* were dominant components of the faba bean mycobiota across sampling periods and plant tissues. Fungal α-diversity and β-diversity were higher during the flowering stage compared with the ripening stage. Basidiomycetous yeasts exhibited greater relative abundance during flowering, whereas *Cladosporium* was significantly more abundant during ripening. Network analysis revealed strong positive associations among Basidiomycota yeasts in June, indicating early-season clustering of yeast taxa within the fungal community. In July, *Alternaria* displayed stronger associations with other members of the order *Pleosporales*, suggesting a mid-season intensification of interactions within *Dothideomycetes*. The α-diversity was significantly higher in asymptomatic plant tissues compared with symptomatic tissues; however, β-diversity was not significantly affected by the presence of disease symptoms. Overall, these findings underscore the importance of precise identification of *Alternaria* species and further investigation of their ecological interactions with faba bean.

## Acknowledgements

This research was funded by Strengthening the Institutional Capacity of LBTU for Excellence in studies and research, funded by the Recovery and resilience facility, grant number Nr. 5.2.1.1.i.0/2/24/I/CFLA/002 and “Research of *Alternaria* spp. and *Stemphylium* spp. as potentially devasting pathogens of faba beans”.

